# TRPM7 regulates phagocytosis and clearance of *Candida albicans*

**DOI:** 10.1101/2023.08.26.554944

**Authors:** Marta E. Stremska, Eric J. Stipes, Jessica J. Jang, Gregory W. Busey, Wesley H. Iobst, Philip V. Seegren, Joel Kennedy, Bimal N. Desai

**Affiliations:** Microbiology, Immunology and Cancer Biology Department, University of Virginia, Pinn Hall, 1340 Jefferson Park Avenue, Charlottesville, VA 22908; Pharmacology Department, University of Virginia, Pinn Hall, 1340 Jefferson Park Avenue, Charlottesville, VA 22908; Carter Immunology Center, University of Virginia, 345 Crispell Dr. MR-6, Charlottesville, VA 22908; Robert M. Berne Cardiovascular Research Center, University of Virginia, 415 Lane Rd, Charlottesville, VA 22908; Department of Immunology and Pathology, Washington University in St. Louis, 425 S Euclid Avenue, St. Louis, MO 63110; Department of Molecular Biology and Genetics, Johns Hopkins Medical School, Baltimore, MD 21205

**Keywords:** Phagocytosis, TRPM7, ion channels, fungi, immunity, ion channels, macrophages

## Abstract

Sentinel phagocytes of the innate immune system have a critical role in detecting and eliminating fungal pathogens. We used patch clamp electrophysiology to explore the electrical signals elicited when macrophages engulf *Candida albicans*. In the perforated patch configuration, which is least disruptive to intracellular signaling, we detected a composite outwardly rectifying current during the engulfment of *C. albicans* or zymosan. FTY720, a known inhibitor of ion channel TRPM7, suppressed the current. We then tested the hypothesis that TRPM7 regulates the engulfment and clearance of *C. albicans*. We found that *Trpm7-/-* macrophages are highly deficient in the engulfment of *C. albicans. Trpm7-/-* macrophages initiate phagocytosis of yeast but are defective in sealing the phagocytic cups. While the precise mechanism through which TRPM7 regulates phagosome sealing is not clear, we tested the immunological significance of this discovery using a mouse model of systemic candidiasis. We show that in mice, wherein TRPM7 is deleted selectively in the myeloid cells, infection by *C. albicans* results in significantly higher lethality, increased colonization of vital organs and increased inflammatory cytokines in the blood. Our study establishes TRPM7 as an ion channel critical for the innate immune responses against fungal pathogens and sets the stage for cell biological studies that define the mechanisms through which TRPM7 regulates phagosome sealing.

**Significance statement:** The worldwide increase in deadly or persistent fungal infections has prompted the research for alternative ways of treatment. We applied the specialized, perforated patch clamp technique to track and identify electrical currents elicited during the detection and engulfment of fungi by macrophages. The ion channel TRPM7 emerged as an important determinant of anti-fungal host defense as its deletion in the murine myeloid cells made the host mice highly susceptible to lethal candidiasis. Ion channels are attractive drug targets whose activation and inhibition can be manipulated with pharmacological therapeutics. This study raises the possibility of enhancing fungal clearance using activators of TRPM7. Such pharmacological strategy may benefit patients of persistent fungal infections that are recalcitrant to anti-fungal drugs.

## INTRODUCTION

Fungal infections pose a major risk to individuals with weakened immune system. *Candida albicans* (*C. albicans*), normally a commensal microbe, can result in persistent infections of mucosal surfaces in immunocompromised patients. Severe candidiasis that is refractory to anti-fungal medications can culminate in colonization of vital organs and sepsis (1). The emergence of drug-resistant strains (2) highlights the urgent need to identify new therapeutic strategies to bolster the defenses against *C. albicans* and related fungal pathogens. Myeloid phagocytes play a crucial role in anti-fungal host defense, and a better understanding of their underlying processes may reveal novel drug targets to promote the clearance of persistent fungal infections.

Phagocytes recognize and engulf fungi through receptors that bind to chemical motifs present in the fungal cell wall (3, 4). The pattern-recognition receptor, Dectin-1 (encoded by *CLEC7A*), is a member of the C-type lectin receptor family that binds to fungal β-1,3 glucans. The proximal downstream signals of the ligand-bound Dectin-1 homodimers involve the phosphorylation of the core tyrosine residue in their hemITAM domain and recruitment of the Syk kinase. Among the downstream Syk substrates, the phosphorylation of phospholipase Cγ (PLCγ) (5-7) is critical for electrical signals, which includes Ca^2+^ signals, that regulate the extension and fusion of pseudopods during phagocytosis (8, 9). The profound changes in membrane phospholipid composition can activate many ion channels but the prevailing framework centers largely on the process of store-operated Ca^2+^ entry (SOCE) and calcium signaling. During SOCE, the activation of IP3 receptors (IP3R) releases the ER-stored Ca^2+^ into the cytosol and subsequent depletion of ER Ca^2+^ stores triggers the activation of plasma membrane resident, Ca^2+^-selective Orai channels. Although SOCE is widely conjectured to be important for phagocytosis (10-12), we are confronted by the paradoxical observation that macrophages deficient in SOCE are able to phagocytose and kill fungi normally (13). We reasoned that electrical signals mediated by other ion channels are more important for the membrane dynamics associated with phagocytosis of fungal pathogens. In this study, we identified TRPM7 as a major player in phagocytosis of fungi and we demonstrate its immunological significance in a mouse model of systemic candidiasis.

TRPM7 is a cation-selective ion channel that belongs to the <u>T</u>ransient <u>R</u>eceptor <u>P</u>otential (TRP) channel superfamily. TRPM7 conducts most cations, including Ca^2+^, Na^+^, Mg^2+^, K^+^, and also possesses a C-terminal serine-threonine kinase domain which has been shown to regulate actomyosin contractility through phosphorylation of Myosin IIA heavy chain (14, 15). Robust TRPM7 currents are readily evident in all hematopoietic cells but in addition to its presence in the plasma membrane, TRPM7 is also emerging as an intracellular ion channel that is located in mysterious vesicles that are not yet well-defined (16). As is typical of TRP channels, TRPM7 can be gated polymodally through variety of intracellular and extracellular signals. TRPM7 can be gated by caspase-mediated cleavage (17), reactive oxygen species (ROS) (16), pH (18, 19), and membrane phospholipids (20). TRPM7 regulates both innate and adaptive immune functions (21-25) and recently, we discovered that in macrophages, the ion channel activity of TRPM7 is critical for the acidification of nascent phagosomes during efferocytosis (26). *Trpm7-/-* macrophages engulf apoptotic cells as well as bacteria (*E*.*coli* was tested) normally. Surprisingly, in this study, we find that *Trpm7-/-* macrophages are highly deficient in the engulfment of *S. cerevisiae and C. albicans. Trpm7-/-* macrophages initiate phagocytosis but fail to seal the nascent phagocytic cups normally. Although the biochemical mechanism through which TRPM7 regulates phagosome sealing remains unclear, the deficit is evidently significant for host defense against fungal pathogens. Mice wherein *Trpm7* is deleted selectively in myeloid cells are highly susceptible to lethal systemic candidiasis and display high fungal loads in their vital organs.

## RESULTS

### In perforated patch configuration, macrophages elicit an outwardly rectifying composite current upon sustained contact with *C. albicans*

To test whether ion channels are activated during phagocytosis of fungal particles, we employed <u>P</u>erforated <u>P</u>atch-<u>C</u>lamp <u>E</u>lectrophysiology (PP-CE), which uses antibiotic-generated pores in the plasma membrane to gain electrical access **(Fig.1A)**. In contrast to Whole Cell Patch-Clamp Electrophysiology (WC-PCE), which gains access by rupturing the cell membrane, PPCE preserves the intracellular environment and signaling, enabling measurements of composite electrical activity in more physiological conditions. However, the inability to control the intracellular ionic composition limits the ability to isolate currents mediated by specific ion channels. Using PPCE, we recorded the composite currents in RAW 264.7 macrophages actively engulfing zymosan particles and observed that the cells develop an outwardly rectifying current **(Fig.1B)**. The summarized current densities show a significant increase in current density in response to zymosan particles **(Fig. 1C)**. Since the steep outward rectification of the current is similar to that of TRPM7, an ion channel that is expressed in macrophages, we tested whether it is sensitive to FTY720, a known TRPM7 inhibitor. The current was inhibited by FTY720 **(Fig.1D)** suggesting that I_TRPM7_ is a significant component of this composite current.

**Figure 1.**
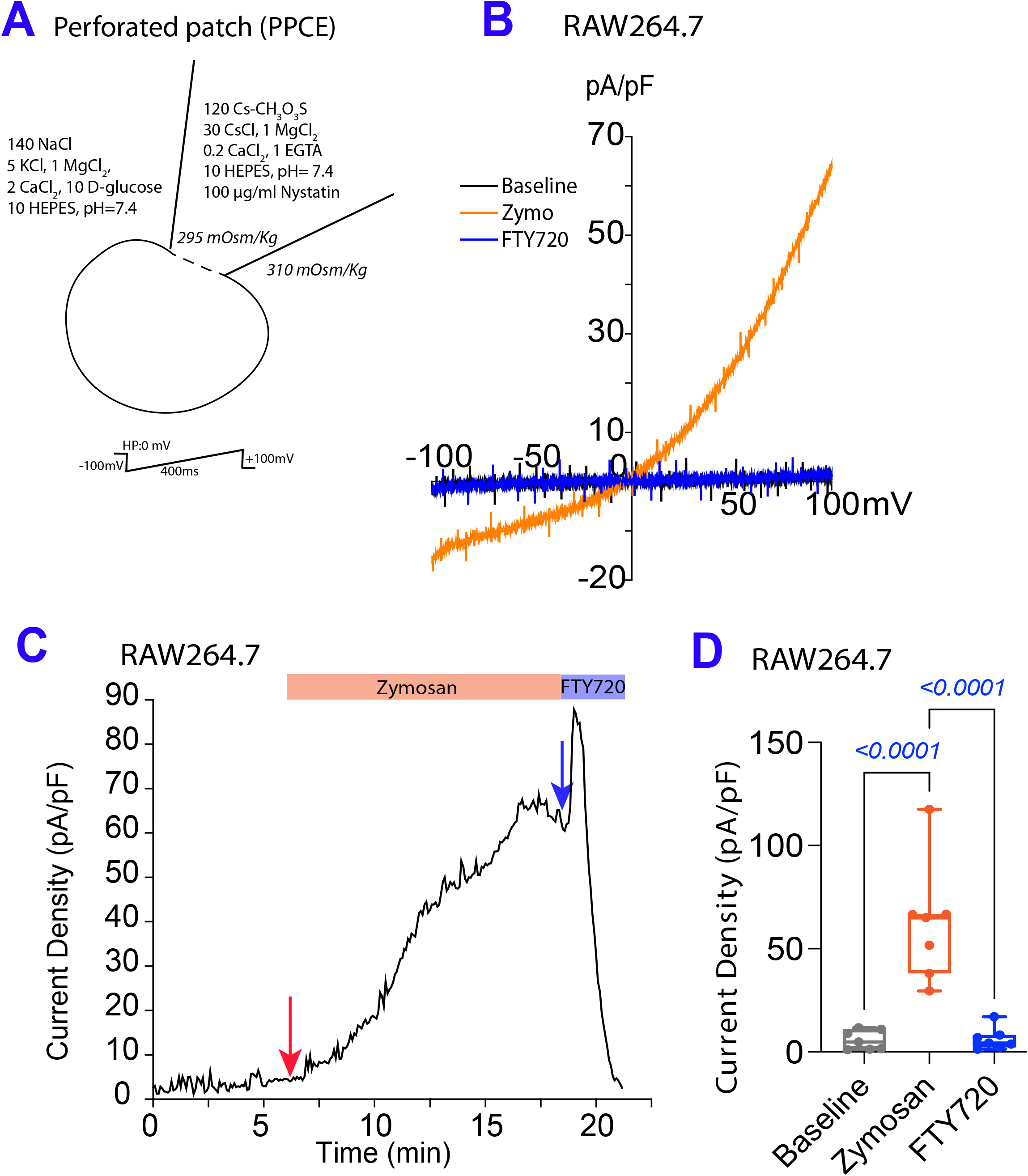
Macrophages elicit an outwardly rectifying composite current in response to fungal particles. (A) Cartoon representation of the conditions in the perforated patch-clamp electrophysiology (PPCE) configuration. The composition of internal (pipet) and external (bath) solutions are outlined but the internal solution does not dialyze the cell through the perforated patch. The voltage ramp is shown together with the filtering and sampling parameters used for signal processing. Holding potential (HP) was at 0mV. (B) Representative I-V relationship of engulfment-triggered current (ETC) in RAW264.7 cells recorded in PPCE configuration; showing baseline trace (black), zymosan-activated current (orange) and FTY720 (5 μM)-inhibited current (blue). Note that the black and blue traces are overlapping. (C) Current densities of ETC obtained in RAW264.7 cells using PPCE; showing baseline (gray, no currents), zymosan-activated currents (orange), and FTY720 (5 μM)-inhibited currents (blue). All currents were quantified at 100 mV. Statistics were performed with ordinary one-way ANOVA. (D) Current density over time in RAW264.7 cells, after the addition of zymosan (red arrow) in a PPCE configuration. Current increased over time and stabilized at ∼10 min. FTY720 (5 μM) inhibited the current. The total recording time was 23 min.

### Maximal TRPM7 currents, isolated using the whole cell configuration (WC-PCE) and Mg^2+^-free conditions, are not different in cells engulfing zymosan

I_TRPM7_ can be isolated more precisely using the whole cell configuration and Mg^2+^-free conditions (27-29), but in these conditions, macrophages do not initiate phagocytosis. Although WC-PCE is not suitable for detecting real-time TRPM7 activation in physiological conditions, it can reveal changes in maximal currents possible between two sets of cells. For instance, if more TRPM7 channels were inserted in the plasma membrane during phagocytosis of fungi, the maximal TRPM7-mediated currents in macrophages that have engulfed yeast would be significantly higher than macrophages that have not been exposed to the fungi. In WC-PCE, we found that I_TRPM7_ density in RAW 264.7 cells fed *C. albicans* for 30 min was not significantly different from current density in unfed cells (Fig. S1B-S1C). Overall, these findings suggest that there is no net change in the number of TRPM7 channels in the plasma membrane during this process.

### *Trpm7-/-* macrophages are defective in engulfment of yeast

To assess the role of TRPM7 in phagocytosis of yeast (*S. cerevisiae* and *C. albicans*), we tested the primary bone marrow-derived macrophages (BMDMs) from *Trpm7*^*fl/fl*^ *LysM-Cre* (KO) mice (26, 30) in *ex vivo* engulfment and killing assays **(Fig. 2A-2B)**. After 30 minutes of exposure to the yeast, the *Trpm7-/-* macrophages engulfed nearly 50% fewer fungi than their *wt* counterparts. This was the case for both *S*.*cerevisiae* and *C. albicans* **(Fig.2C)**. However, we saw no difference in the killing of the engulfed yeast particles **(Fig. 2B)**. We also used transmission electron microscopy (TEM) to visualize yeast-containing phagosomes **(Fig. 2D)** and counted the number of internalized fungi in primary macrophages. Consistently the *Trpm7-/-* macrophages showed fewer internalized yeast **(Fig. 2E)**. We considered the possibility that the engulfment deficit is a secondary result of a primary deficit in macrophage migration, which would prevent the macrophages from seeking the yeast in the cell cultures. However, when we assessed the migratory ability of *Trpm7-/-* macrophages using a Boyden chamber (31) (*outlined in* Fig.S1B), we found no significant defects in their migration toward *C. albicans* (Figure S2B-S2C). The cell viability of *Trpm7-/-* macrophages cultured with the yeast was also comparable to that of the *wt* macrophages, indicating that they were not dying in abnormally high numbers in response to yeast pathogens (Fig. S2D). Phagocytes use several pattern-recognition receptors (PRR) to recognize and engulf fungal pathogens. These receptors include Toll-like receptors (TLRs) (32, 33), C-type lectin receptors (CLRs) (34, 35), NOD-like receptors (NLRs) (36), and RIG-I-like receptors (RLRs) (37). We found no significant differences in the expression levels of these PRR between *wt* and *Trpm7-/-* macrophages (Fig. S2E). Overall, these findings indicate that TRPM7 regulates phagocytosis of fungi. Additionally, it is clear that the engulfment defect in *Trpm7-/-* macrophages is not due to deficits in migration, viability or reduced expression of key receptors.

**Figure 2.**
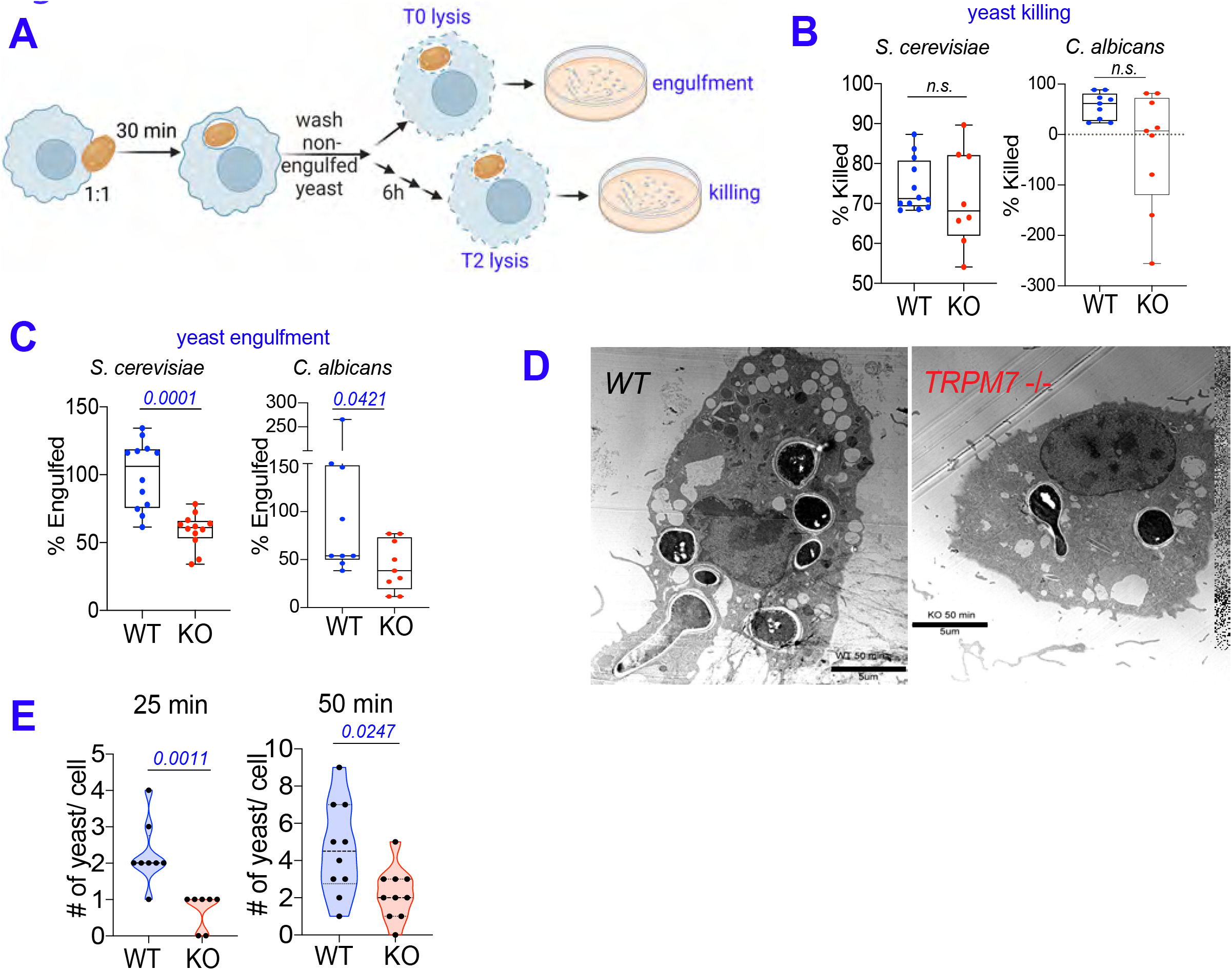
*Trpm7-/-* macrophages are defective in the engulfment of yeast. (A) A cartoon schematic outline of the *in vitro* yeast killing assay. (B) The yeast killing efficiency of WT (blue), and KO (red) BMDMs, as quantified for *S. cerevisiae* (*left panel*) and *C. albicans* (*right panel*) engulfment assays. Statistics were performed with a two-tailed *t* test. (C) Quantification of yeast engulfment of WT (blue), and KO (red) BMDMs, as quantified for *S. cerevisiae* (*left panel*) and *C. albicans* (*right panel*). Statistics were performed with a two-tailed *t* test. (D) Representative transmission electron microscope (TEM) micrographs for WT (*left*) and KO (*TRPM7-/-, right*) macrophages at 50 min post *C. albicans* engulfment. Scale bar= 5 μm. (E) Quantification of engulfment of *C. albicans* per cell, derived from TEM micrographs at two tested time points: 25 min (*left*) and 50 min (*right*) for WT (blue shaded violin plot, 25 min *n*=8; 50 min *n*=10) and KO (red shaded violin plot, 25 min *n*=7; 50 min *n*=10). Statistics represent a two-tailed *t* test.

### TRPM7 regulates the sealing of the nascent yeast-containing phagocytic cups

To investigate the possible role of TRPM7 in the earliest stages of phagocytosis, we assessed the formation and sealing of nascent phagosomes containing the yeast. In this experiment, we used yeast particles that are cell surface biotinylated and labeled with Streptavidin-568 fluorophore (red fluorescent). As illustrated **(Fig. 3A)**, this enables the visualization of nascent phagosomes containing the yeast. Importantly, when the nascent phagosome is successfully sealed, the yeast is no longer accessible to labeling by the second streptavidin-conjugated fluorophore (Streptavidin-488, green fluorescent). We offered the modified yeast to the *wt* and *Trpm7-/-* macrophages for engulfment. After 15 and 30 minutes, the unattached or loosely attached yeast were washed off and the macrophages engulfing the yeast were labeled with Streptavidin-488, fixed, and the preparations visualized by fluorescence confocal microscopy. Through such analyses, we visualized and quantified the efficiency of phagosome formation and sealing in *wt* and *Trpm7-/-* macrophages. These results **(Fig. 3B)** reveal that after 30 minutes of engulfment, the wt macrophages are successful in sealing their phagosomes – the engulfed yeasts are labeled with Streptavidin-568 but not with Streptavidin-488. However, the *Trpm7-/-* macrophages display many unsealed phagosomes – the yeast in these unsealed phagosomes is labeled with both Streptavidin-568 and Streptavidin-488. Quantifying these differences across many confocal images shows that the *Trpm7-/-* macrophages are significantly less efficient at sealing the phagosomes. The percentage of fully engulfed yeast in *Trpm7-/-* macrophages was significantly lower **(Fig. 3C)** and accordingly, the percentage of non-engulfed (includes partially engulfed) yeast was significantly higher **(Fig. 3D)**. However, these differences were not evident at 15 minutes past engulfment because at this time point, the formation of phagosomes in both *wt* and *Trpm7-/-* macrophages was comparable and these phagosomes were unsealed in both macrophage populations (Fig. S3A). Overall, the results indicate that TRPM7 regulates the sealing of nascent phagosomes.

**Figure 3.**
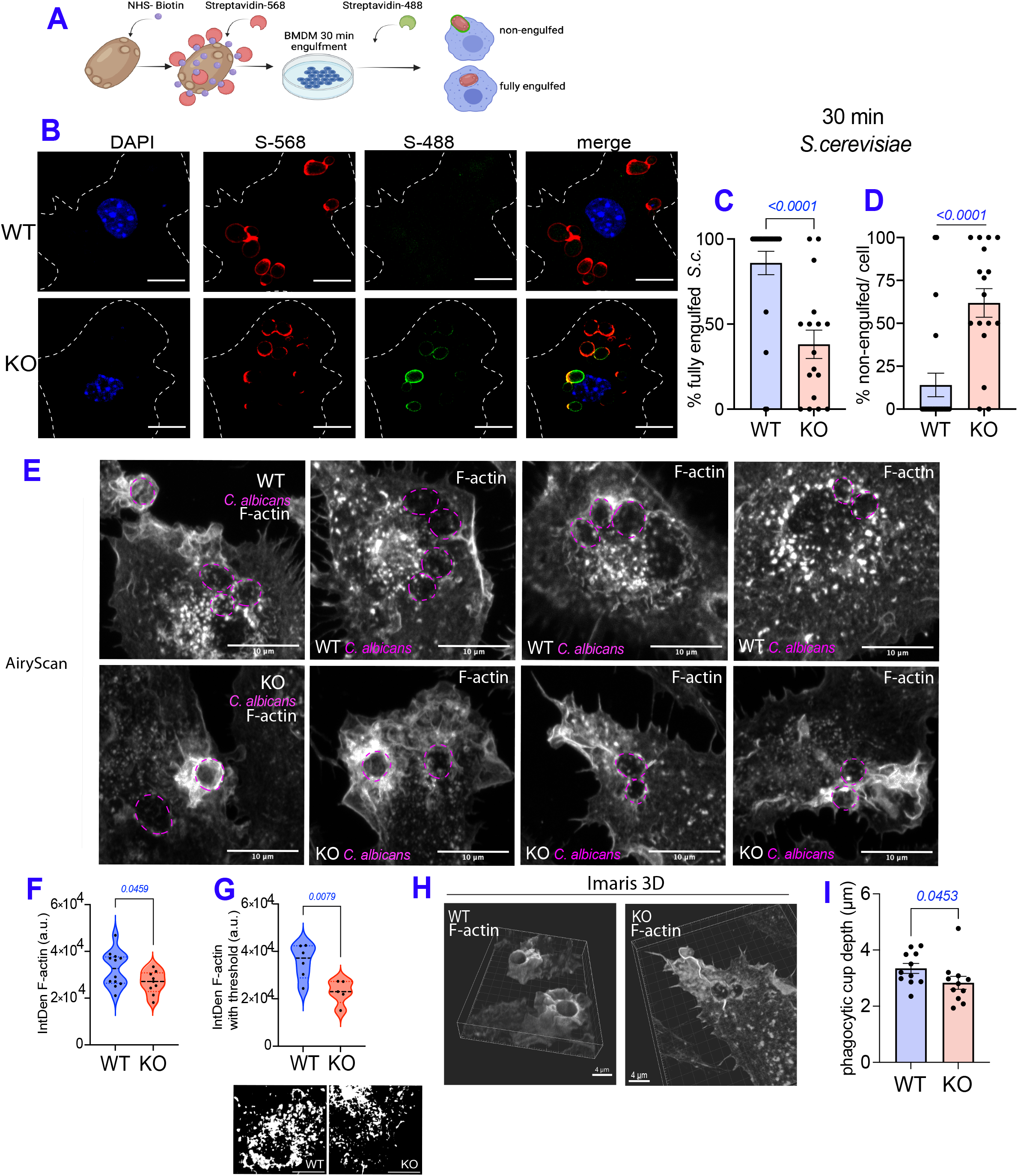
TRPM7 regulates the sealing of the nascent yeast-containing phagocytic cups. (A) A cartoon showing the experimental design to evaluate the phagosome sealing during yeast engulfment. (B) Representative confocal images of *S. cerevisiae* engulfment at 30 min, for WT (*upper panel*) and KO (*lower panel*) macrophages. Single channels and merged images are shown. Scale bar=10 μm. (C) Quantification of (B) showing fully engulfed *S. cerevisiae*, expressed as a percentage at 30 min engulfment. Statistics show a two-tailed *t* test with SEM. (D) Quantification of (B) showing the percentage of partially engulfed (unsealed phagosomes) *S. cerevisiae* at 30 min engulfment. Statistics: two-tailed *t* test with SEM. (E) Maximal z-stack projections of F-actin staining in WT (*upper panels*) and KO (*lower panels*) macrophages at 5 min incubation with *C. albicans* (*outlined in magenta-dashed circles*). Scale bar=10 μm. Representative images of four different cells for each genotype. (F) Quantification of the integrated density (IntDen) of F-actin staining in WT (*blue*) and KO (*red*) primary macrophages. Statistics: two-tailed *t* test with SEM. (G) Quantification of the integrated density (IntDen) after applying the threshold that isolated the F-actin-rich puncta. Statistics: two-tailed *t* test with SEM. (H) Imaris 3D rendition of z-stacks acquired from Airy Scan confocal microscopy, showing phagocytic cup formation in WT and KO macrophages after 5 min incubation with *C. albicans*. Scale bars are outlined in each image. (I) Quantification of the phagocytic cup depth in WT (*blue*) and KO (*red*) BMDMs. Statistics show a two-tailed *t* test.

### The phagocytic cups in Trpm7-/-macrophages show morphological differences

A recent study showed that F-actin nucleation exerts constricting forces around the phagocytosed particle while myosin II forms contractile rings around the phagocytic cup, which are crucial for its closure (38). Since TRPM7 is known to regulate actomyosin contractility and its kinase domain can phosphorylate myosin IIA heavy chain (14, 15), we sought to visualize F-actin nucleation in in the earliest stages of phagocytosis. Confocal imaging of live cells labeled with Actin tracking fluorescent dye (CellMask) revealed that wt macrophages phagocytosing *C. albicans* formed dense, actin-rich podosome structures **(Fig.3E)**, which are thought to facilitate phagocytosis by serving as initiation sites for actin polymerization (39, 40). The *Trpm7-/-* macrophages formed significantly fewer such podosome-like structures **(Fig. 3F)** and this was especially evident if the quantification was restricted to structures that achieved an arbitrary threshold of fluorescence density **(Fig. 3G)**. To assess differences in the phagocytic cups containing the engulfed yeast, we used the z-stacks of confocal images and rendered them as 3D projections that are amenable to measurements **(Fig.3H)**. The phagocytic cups in *Trpm7-/-* macrophages displayed reduced depth, suggesting that the progression of phagocytosis was slower **(Fig. 3I)**. Using immunofluorescence confocal microscopy, we also found that in *Trpm7-/-* macrophages, the distribution of myosin IIA was different, displaying relatively more filamentous structures (Fig.S3B). However, total levels of myosin IIA and myosin 9 were not different in *Trpm7-/-* macrophages (Fig. S3C). Overall, these morphological differences substantiate the idea that TRPM7 regulates the cytoskeletal dynamics during phagocytosis of *C. albicans*.

### Increased lethality from candidiasis in mice wherein Trpm7 is deleted selectively in myeloid cells

To assess whether the role of TRPM7 in phagocytosis of *C. albicans* was physiologically significant for host defense, we used a mouse model of systemic candidiasis. The mice were administered *C. albicans* via intravenous (i.v.) injection and monitored for survival and severity of disease (41, 42). As shown in the Kaplan-Meier survival curve analysis we found that when compared to *wt* mice, candidiasis was strikingly more lethal in *TRPM7*^*fl/fl*^ *LysM-Cre* (KO) mice (n=10, p=0.0081), with a median survival time of 73h **(Fig. 4A)**. In a modified experiment where the mice were infected with a smaller dose of *C. albicans*, the KO mice also lost more weight than their *wt* counterparts, over 48h of infection **(Fig. 4B)**. To assess the colonization of organs by *C. albicans*, we used PAS histochemical stain, which detects the fungal polysaccharides in tissue sections. PAS staining of kidney sections, derived 48h after infection, revealed larger and far more numerous yeast colonies throughout the renal cortex and outer medullary regions in the KO mice, when compared to *wt* mice **(Fig. 4C)**. When these colonies quantified in terms of area, the pathogen burden was significantly higher in the KO mice **(Fig. 4D)**. We used immunohistochemistry to assess immune infiltration in the infected kidneys. Infiltration by CD45+ hematopoietic cells was evident in both *wt* and *KO* kidney sections (Fig. S4C) and when quantified, there were no statistically significant differences (Fig. S4D). However, there were significantly more neutrophils (Ly-6G+) in kidney sections of the KO mice **(Fig. 4E)**. Analysis of serum inflammatory cytokines showed signs of modestly increased inflammation in the infected *Trpm7*^*fl/fl*^ *(LysM Cre)* mice, when compared to their *wt* counterparts. G-CSF, in particular, was significantly elevated in the *Trpm7*^*fl/fl*^ *(LysM Cre)* mice **(Fig. 4G)** and this may be an important contributor to the increased infiltration of neutrophils in the kidneys (43). We also assessed the overall pathogen load by plating tissue homogenates on YPD-agar and quantifying the colony forming units (CFUs) per gram of tissue. In this assay, the liver and kidney homogenates from KO mice yielded more CFUs but the difference was not statistically significant (Fig. S4A and Fig. S4B). The findings from the mouse model of systemic candidiasis lead to the conclusion that TRPM7 function in myeloid phagocytes is crucial for host defense against infection by *C. albicans*.

**Figure 4.**
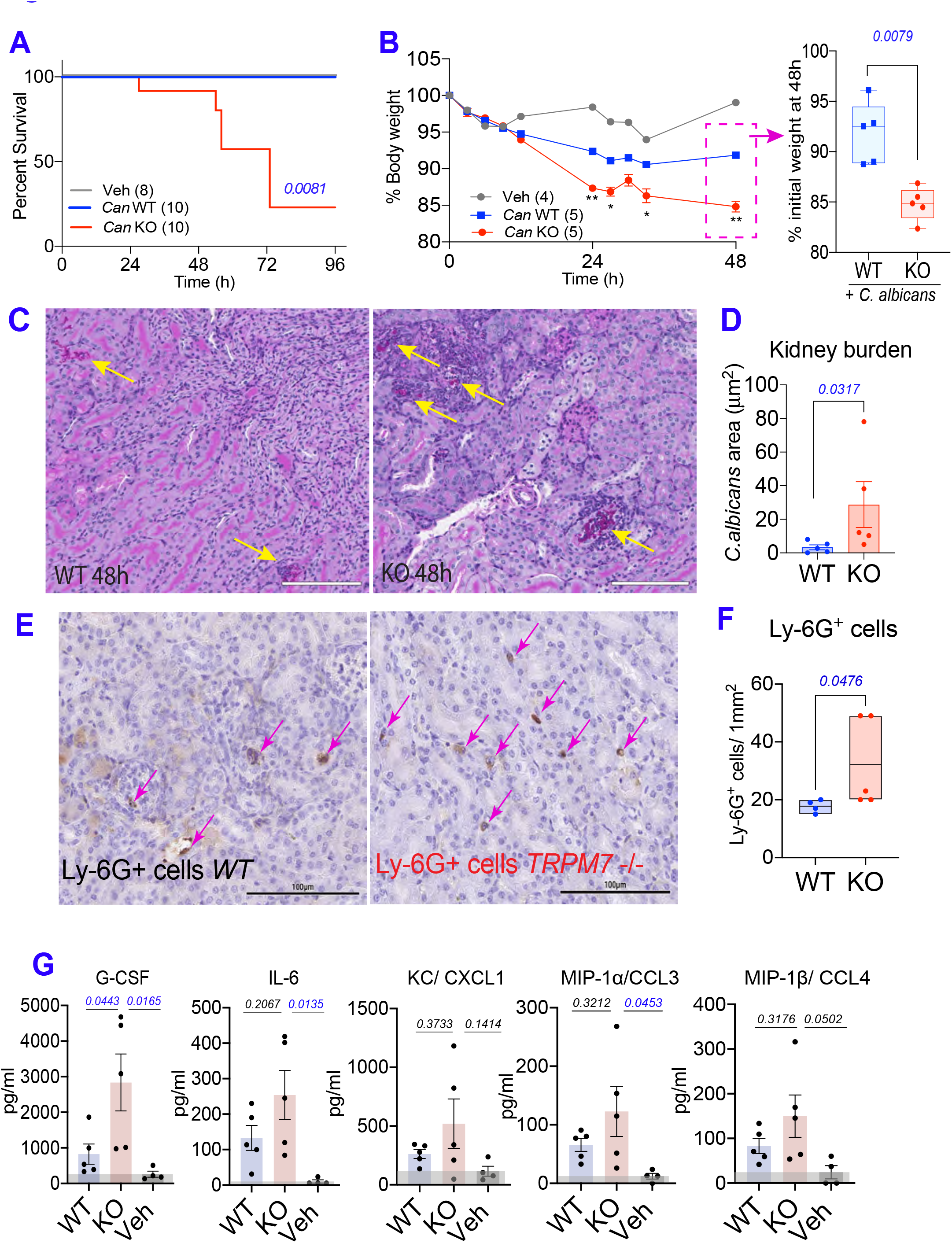
*Trpm*^*fl/fl*^ *LysM-Cre* mice show increased susceptibility to lethal candidiasis. (A) Kaplan-Meier survival analysis of *TRPM7*^*fl/fl*^ (WT; blue, *n*= 10) and *TRPM7*^*fl/fl*^ *LysM-Cre* (KO; red, *n*=10) mice injected intravenously with 100μl 1x10^6^ *C. albicans* and assessed over 96 hours. Control mice were injected with 0.9% saline solution (Veh) and were both WT (*n*=4) and KO (*n*=4) outlined in a grey line. Curve comparison was performed using a log-rank (Mantel-Cox) test with a *p*= 0.0081 (B) Body weight loss over time, presented as a % of initial weight for WT (*n*=5, blue), KO (*n*=5, red), and vehicle-injected control mice (WT, *n*=2; KO, *n*=2). Statistics performed with a 2way AVOVA with multiple comparisons \**p*=<0.05, ** *p*=<0.005. *Right panel*: box and whisker plot (min to max) showing cumulative % initial body weight at 48 h. Statistics were performed with a two-tailed *t* test (Mann-Whitney) (C) Periodic-acid Schiff (PAS) stained kidney sections, collected at 48 h from WT and KO mice (*n*=5 each; WT-blue, KO-red) injected with *C. albicans*. Yellow arrows indicate *C. albicans* colonies (D) Quantification of PAS staining by outlining and counting the affected areas (in μm^2^) colonized by *C. albicans* in each kidney section. Statistics performed with a two-tailed *t* test (Mann-Whitney) with SEM (E) Representative kidney sections stained with anti-Ly-6G antibody harvested from WT (*left panel*) and KO (*right panel*) mice at 48h. Magenta arrows indicate positively stained cells (brown color). The scale bar is 100 μm. (F) Quantification of anti-Ly-6G stained kidney sections per 1 mm^2^ area, 48 h post *C. albicans* injections in WT (blue, *n*=5) and KO (red, *n*= 5) mice. Data were plotted as floating bars and statistics were performed using two-tailed *t* test with SEM. Measurement of serum cytokines at 48h post-*C. albicans* injections via Luminex Multiplex assay. The data were plotted for WT (*n*= 5, blue bars), KO (*n*=5, red bars), and PBS controls (grey bars, *n*=4). Statistics performed with an ordinary one-way ANOVA, error bars represent SEM

## DISCUSSION

TRPM7 is emerging as an important ion channel for innate immunity and macrophage biology. In our previous studies on the function of TRPM7 in macrophages, we discovered that TRPM7 is not required for the engulfment of apoptotic cell corpses but is critical for the maturation of the phagosomes during efferocytosis. *Trpm7-/-* macrophages are normal in phagocytosis and killing of *E*.*coli* but TRPM7 regulates LPS-triggered TLR4 signaling and macrophage activation. In the context of these previous discoveries, the newly identified function of TRPM7 in the engulfment of *C. albicans* suggests a framework in which TRPM7 regulation and function is contextual, responding to the subtle variations in upstream signals.

We show evidence indicating that TRPM7 regulates the sealing of nascent phagocytic cups. TRPM7 is known to regulate cytoskeletal rearrangements and actomyosin contractility (14, 15, 44). We speculate that TRPM7 regulates the sealing of phagocytic cups through the regulation of myosin IIA contractile ring around the phagocytic cup. Alternatively, or additionally, TRPM7 may regulate the formation of actin-dense protrusions called podosomes (15, 45). *In vivo*, podosomes enable macrophages to invade through the extracellular matrix (46-48). This aspect could be critical for the phagocytic clearance of fungal pathogens in the complex 3D tissue architecture. On a molecular scale, TRPM7 is likely to regulate these processes through the combined activities of its ion channel and kinase domains. The TRPM7 kinase can phosphorylate Myosin IIA heavy chain but there are likely to be other substrates of relevance to engulfment.

The TRPM7 pore is non-selective and can conduct most cations present in physiological fluids. While it is tempting to implicate TRPM7-mediated Ca^2+^-signaling at the site of membrane sealing, the exploration of this aspect poses significant technical challenges. Recognition of fungal β-glucans by Dectin-1 elicits global Ca^2+^ elevations through SOCE. In this context, resolving small and rapid local changes in the subcellular space of the extending pseudopods proved difficult and inconclusive using conventional Ca^2+^ indicators. It is also possible that TRPM7 mediated changes in local membrane potential are important for the extension of pseudopods and sealing of the nascent phagosome. Patch clamp electrophysiology of macrophages yields robust TRPM7 currents so the presence of TRPM7 on the plasma membrane is not in doubt. However, there is increasing evidence that the bulk of the TRPM7 is relatively undefined intracellular vesicles (16). Understanding the trafficking and content of these vesicles may provide new insights into the cell biology of TRPM7 and thereby bring us a step closer to a coherent overarching framework of TRPM7 regulation and function.

Our failure to define a clear molecular mechanism underlying the immunologically significant role of TRPM7 in the phagocytosis of *C. albicans* is clearly a limitation of this study. Deciphering this mechanism will require generation of new tools and reagents and will be a topic of our future research. Overall, despite these limitations, this study provides conclusive evidence that TRPM7 regulates the phagocytosis of *C. albicans* and that its function in myeloid cells is a critical for host defense against *C. albicans*. To the best of our knowledge, TRPM7 is the first and only ion channel shown to be critical for anti-fungal host defense. Since ion channels are attractive drug targets, and can be modulated positively (activated) or negatively (inhibited) through pharmacological means, our study raises the possibility of enhancing fungal clearance using activators of TRPM7. Such a pharmacological strategy may be of clinical benefit in case of persistent fungal infections that are recalcitrant to anti-fungal drugs.

## METHODS

### Mouse Strains

*Trpm7*^*fl/fl*^ and *Trpm7*^*fl/fl*^ *(LysM Cre)* mice were generated as described previously (24) and crossed to B6 background. Male and female mice 8-12 weeks of age were used for all experiments. Deletion of *Trpm7* was confirmed via quantitative real-time PCR analysis and patch-clamp electrophysiology. All animals were bred and housed in line with the policies of the University of Virginia Institutional Animal Care and Use Committee (IACUC).

### Mouse genotyping

Mice were tail clipped upon weaning and the tail samples were dissolved in DirectPCR Lysis Reagent (Viagen Biotech; #102-T), as outlined in the manufacturer’s protocol. 1-2 μl of the crude lysate was used as the PCR template, performed with GoTaq Green (Promega; #M7122).

### *In vivo* disseminated *Candidiasis* mouse model

Age-matched male and female mice were injected i.v. (tail vein) with 1x10^6^ previously washed (3x in 1xHBSS) and counted *C. albicans*, suspended in 0.9% saline solution. Mice were randomly assigned to experimental cages and allocated the appropriate injection by another experimenter. Mice were weighed before and at regular time points post-injection, as outlined in the figure panels and assessed for signs of distress (conjunctivitis, dull fur, hunching, nose bulge). After general anesthesia with i.p.-administered Avertin (2, 2, 2-tribromoethanol, Sigma-Aldrich #T48402) the blood was collected with heparinized capillaries (Fisher Scientific™ #22-260950) and mice were euthanized.

### Cell lines and cell culture

RAW267.4 (ATCC® TIB-71™) cell line was maintained according to the ATCC guidelines, in a humified incubator at 37°C and 5% CO_2_. Bone marrow-derived macrophages (BMDMs) were isolated and cultured as previously described (42). In brief, bone marrow was extracted from the murine femur and tibia via centrifugation. Red blood cells (RBCs) were lysed in ACK Lysis buffer and plated on petri dishes at 2-4x 10^6^ cells/ plate in BMDM Media (RPMI 1640 + 10% FBS + 20% L929-conditioned media). Cells were differentiated for 7 days, and the media was refreshed afterwards every 3 days. For experiments, BMDMs were used between days 7 to 10 after harvest.

### Preparation of yeast cultures

*Saccharomyces Cerevisiae* (strain sy1022 fy5) was provided as a gift from Jeff Smith Lab, UVA, and *Candida Albicans* was purchased from ThermoFisher Scientific #R4601503. Yeast strains were streaked on YPD agar plates. When preparing cultures, a single colony was inoculated into previously autoclaved 5ml YPD broth in a glass tube and grown overnight. The cultures were washed 3x in sterile 1xHBSS and counted on a hemacytometer or based on their OD600, where OD600 of 1 equals 3x10^7^ yeast/ml. The yeast cultures were freshly grown for experiments and kept in the fridge.

### Yeast engulfment and killing assays

Killing assays were performed as described previously (42). Briefly, primary macrophages or RAW264.7 cells were incubated with yeast strains *C. albicans* or *S. cerevisiae* for 30 min at 37°C (5% CO2). Cells were then washed 3x with HBSS to remove non-engulfed yeast and resuspended in BMDM media. One well per sample was harvested and lysed at 30 min time point (T0) in sterile, distilled water for 30 min with shaking. Each well was serially diluted and plated on YPD agar plates. The T2 time point wells were harvested at the indicated times, usually at 6h, and lysed as before. The % yeast killing was calculated by the formula 100%-(T2/T0)x100. For engulfment assays, only T0 samples were used with appropriate treatments. For the biotinylation of *S. cerevisiae* two protocols were adapted (49, 50). Briefly, *S. cerevisiae* was subjected to biotinylation in 1 mg/ml EZ-Link™ Sulfo-NHS-SS-Biotin (Thermo Scientific #21331) for 1h at RT with rotation. Streptavidin conjugated to either Alexa Fluor™ -488 (ThemoFisher #S32354) or -568 (ThermoFisher #S11226) was added to triple-washed biotinylated yeast and again incubated for 1h at RT in dark with rotation.

### Transwell migration assay

The transwell polycarbonate membrane cell culture inserts (#CLS3422; Corning) were used. *C. albicans* were prepared from a fresh overnight culture, as outlined above, and placed in the lower chamber compartment suspended in BMDM media. BMDMS were freshly seeded on the membrane, allowed to adhere for 30 min, and then incubated for 6h. Control samples for basal migration assessment had no *C. albicans* present in the lower chamber. At the end of the experiment, the non-migrated cells, remaining on the top of the membrane were gently scraped off with a clean cotton swab, the membranes were rinsed with PBS and fixed (4% PFA in PBS), stained, and mounted on glass slides using ProLong™ Gold antifade reagent with DAPI (#P36935; Invitrogen). Migration was quantified using light microscope at 40x magnification. The 10 ROIs were chosen randomly and averaged for each condition.

### Phagosomal sealing experiments

For bead biotinylation, a previously published protocol was adapted (50). Briefly, 3 μm (#19118-2; Polysciences) or 6 μm (#17145-4; Polysciences) amine-conjugated beads were washed in anhydrous DMSO (#276855; Sigma-Aldrich) using 0.45 μm PTFE membrane columns (#UFC30LH25; Millipore). Amine groups were activated by 0.5 M CDI (#115533; Sigma-Aldrich) in DMSO and the beads were incubated on a bench-top rotor for 1h at RT. 10 mM β-glucan (#346210; EMD Millipore) was dissolved in DMSO and added to the beads for further 1h incubation at RT (with rotation). Beads were washed using the PTFE filters and resuspended in 1x PBS or further biotinylated with 6 mg/ml EZ-link™ Sulfo-NHS-SS-biotin (#21331; ThermoFisher). Streptavidin conjugated to either AlexaFluor-488 or -568 was then added for 1h RT incubation, as before. The phenol-sulfuric acid assay was performed to validate the conjugation of β-glucan and measure the carbohydrate content. Further validation via RT-qPCR measurement of IL-1β transcript levels was performed on RAW267.4 cells incubated with a 1:1 ratio of 3 μm-β-glucan conjugated beads for 3h.

### Patch Clamp Electrophysiology

All electrophysiology experiments were conducted at RT with an Axopatch 200B amplifier (Molecular Devices, Sunnyvale, CA). To measure TRPM7 currents (I_TRPM7_) in RAW264.7 cells (WC-PCE configuration), macrophages were freshly seeded onto the glass coverslips and allowed to adhere. The composition of the WC-PCE extracellular bath solution was (in mM): 140 Na-CH_3_O_3_S, 5 Cs-gluconate, 2.5 CaCl_2_, 10 HEPES, (adjusted to pH 7.4 with osmolality 285 mOsm/kg). The pipette solution contained (in mM): 115 Cs-gluconate, 3 NaCl, 0.75 CaCl_2_, 1.8 Cs_4_-BAPTA, 2 Na_2_ATP, 10 HEDTA, 10 HEPES (adjusted to pH 7.4 with osmolality 273 mOsm/kg). Inhibition of I_TRPM7_ was achieved by using FTY720 (5 μM). For perforated-patch (PPCE) configuration 100 μg/ml of freshly prepared Nystatin was added in the pipette. Extracellular bath contained (in mM): 140 NaCl, 5 KCl, 1 MgCl_2_, 2 CaCl_2_, 10 D-glucose, 10 HEPES (adjusted to pH 7.4 with osmolality 310 mOsm/kg). The PPCE pipette solution contained (in mM): 120 Cs-methanesulfonate (Cs-CH_3_O_3_S), 30 CsCl, 1 MgCl_2_, 0.2 CaCl_2_, 1 EGTA, 10 HEPES, 100 μg/ml Nystatin (pH= 7.4). Recordings were obtained with holding potential of 0 mV, employing a voltage ramp from -100 mV to +100 mV for 400 ms; filtering the signaling at 5 kHz and sampling at 10 kHz. Zymosan particles and *S. cerevisiae* extract were prepared in PPCE bath solution.

### Immunocytochemistry

Cells were seeded on glass coverslips prior to experiments. Coverslips were washed in 1xPBS prior to fixing with 4% PFA (in PBS) for 20 min at RT and washed 3x in 1xPBS prior to blocking at RT for 1h in blocking buffer (5% donkey serum, 1% BSA, 0.1% fish gelatin, 0.1% Triton X-100, and 0.05% Tween-20 in PBS) or BlockAid™ Blocking Solution (#B10710). Incubation with primary antibodies was performed overnight at 4°C, and they were diluted in a blocking solution at concentrations indicated by the manufacturer. Post-incubation, samples were washed 3x in 1xPBS and incubated with a secondary antibody, conjugated to a fluorophore, for 90 min at RT (in the dark), followed by 3x wash in 1xPBS. For F-actin staining, cells were prepared with CellMask™ Actin (Invitrogen; #A57243 or #A57245), as recommended by the manufacturer. Coverslips were then mounted on glass slides using ProLong™ Gold Antifade #P36930, allowed to cure overnight, and imaged within 48 hours. Confocal microscopy imaging was performed with Zeiss LSM880 with AiryScan and analyzed using Fiji (51).

### Western blot

Whole-cell lysates were prepared by harvesting and incubating cells for 30 min on ice, in the desired volume of Lysis Buffer (300mM NaCl, 1% NP-4, 50mM Tris-HCl, 0.5% sodium deoxycholate, 0.1% SDS, pH= 7.4). Soluble proteins were harvested by centrifugation at 20,000 xg for 15 min in a tabletop centrifuge (4°C). Total protein concentration was assessed by a BCA Assay (ThermoFisher; #23225). Supernatants were mixed with 5X Laemmli Buffer (0.3M Tris-H-Cl, 10% SDS, 50% glycerol, 25% β-mercaptoethanol, 0.05% bromophenol-blue) and boiled at 95°C for 10 min. Samples were loaded onto 4–20% Mini-PROTEAN® TGX™ Precast Protein Gels (BioRad; #4561096) and separated by electrophoresis (150V, 90 min in SDS-PAGE buffer). Protein transfer onto a PVDF membrane was performed with a Trans-Blot Turbo Transfer system (BioRad).

### qRT-PCR gene expression

Macrophages were seeded onto a 12-well plate at a density of 0.5x 10^6^ and cultured overnight. The next day they underwent stimulation as outlined in the figure panels. When using pharmacological treatment, the appropriate drug was added 10 min prior to stimulation with fungal particles in complete media and then together with the fungal particles. RNA isolation was performed using RNeasy Plus Kit (Qiagen#74134) and cDNA synthesis was performed using GoScript™ Reverse Transcriptase Kit (Promega #A5001). qRT-PCR reactions were performed using SensiFast SYBR no-rox kit (BIO-98020).

### Tissue processing and immunohistochemistry

Mouse tissues processing, H&E and PAS staining were performed at the University of Virginia Research Histology Core (Sheri VanHoose, MLT [NCA]). The Immunocytochemistry with CD45 (#550539, BD Biosciences) and Ly-6G (#12762, Biolegend) antibodies, together with slide scanning were performed at the University of Virginia Biorepository and Tissue Research Facility (BTRF; Pat Pramoonjago, PhD).

### Statistics

The data analysis was performed with Excel (Microsoft), GraphPad Prism 9.0 (GraphPad Software), and Origin Pro 9.1.0 (Origin Lab), plotted with GraphPad and Origin Pro. Data was preferably presented as individual data points for each independent sample and plotted with means and error bars, as described in figure legends. The sample size and p values are indicated in figure legends, *p* values less than 0.05 were considered statistically significant.

## DATA AVAILABILITY STATEMENT

All data, unique reagents and code we generated for this manuscript will be available upon request to the corresponding author (BND).

## AUTHOR CONTRIBUTIONS

**Conception:** MES, BND; **Research Design:** MES, BND; **Experimental execution:** MES, JJJ, EJS, WHI, GWB, PVS; **Data analysis:** MES, EJS, JJJ, GWB, WHI, PVS; **Technical assistance:** EJS, JK; **Manuscript writing:** MES, BND; **Project Administration:** BND

The authors declare that they have no conflicts of interests.

## ACKNOWLEDGEMENTS

We would like to acknowledge members of the Desai Lab for helpful comments and suggestions. We also thank the members of the Ravichandran Lab (UVA) for technical assistance and feedback. We used the following UVA core facilities: Advanced Microscopy Facility, Flow Cytometry Core, Research Histology Core and Biorepository and Tissue Research Facility for their expertise and support in this project.

This work was supported by NIGMS grant to B.N.D. (GM108989)

---

**Supplementary Figure 1.**
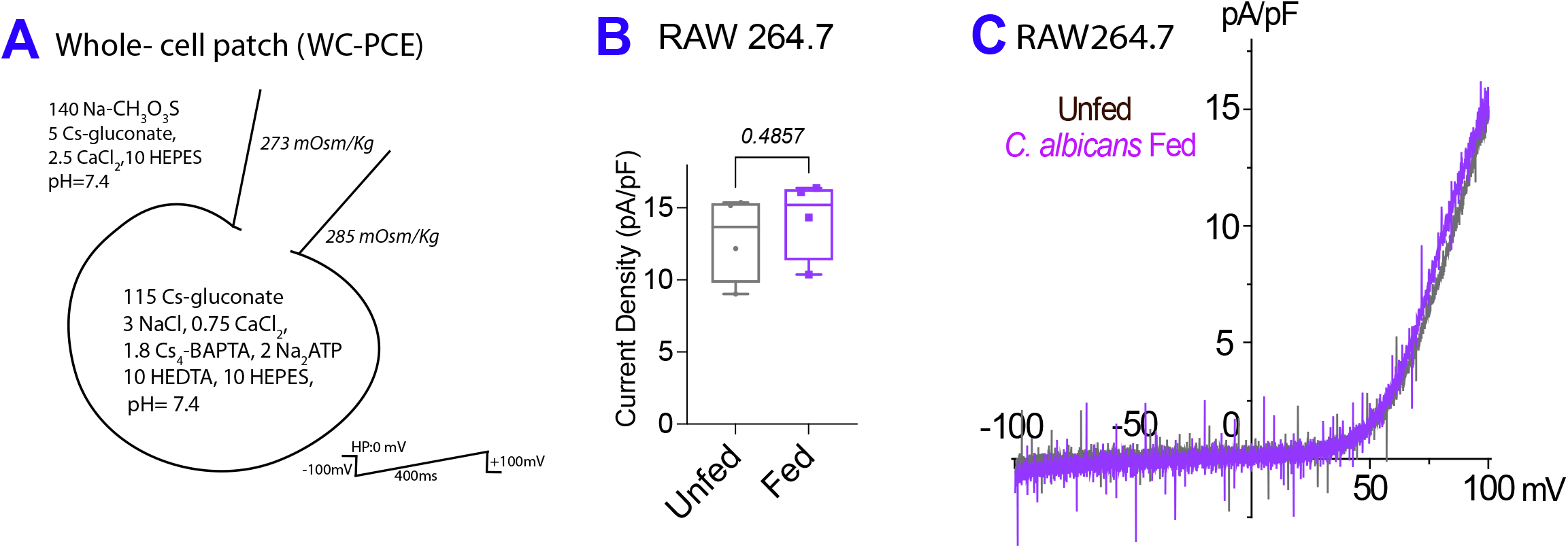
(A) Cartoon of the conditions in Whole-cell patch-clamp electrophysiology (WC-PCE) configuration. The composition of internal (pipet) and external (bath) solutions are outlined. The voltage ramp is shown together with the filtering and sampling parameters used for signal processing. HP was at 0mV. (B) TRPM7 current densities in RAW cells in WC-PCE configuration, before and 30 min post-incubation with *C. albicans* (Fed). Currents were quantified 5 min after break-in at 100 mV. Statistics were performed with a two-tailed *t* test with SEM. (C) Representative current-voltage (I-V) relationship of I_TRPM7_ in untreated (black trace) and *C. albicans* treated RAW cells (30 min, purple trace) obtained in WC-PCE configuration.

**Supplementary Figure 2.**
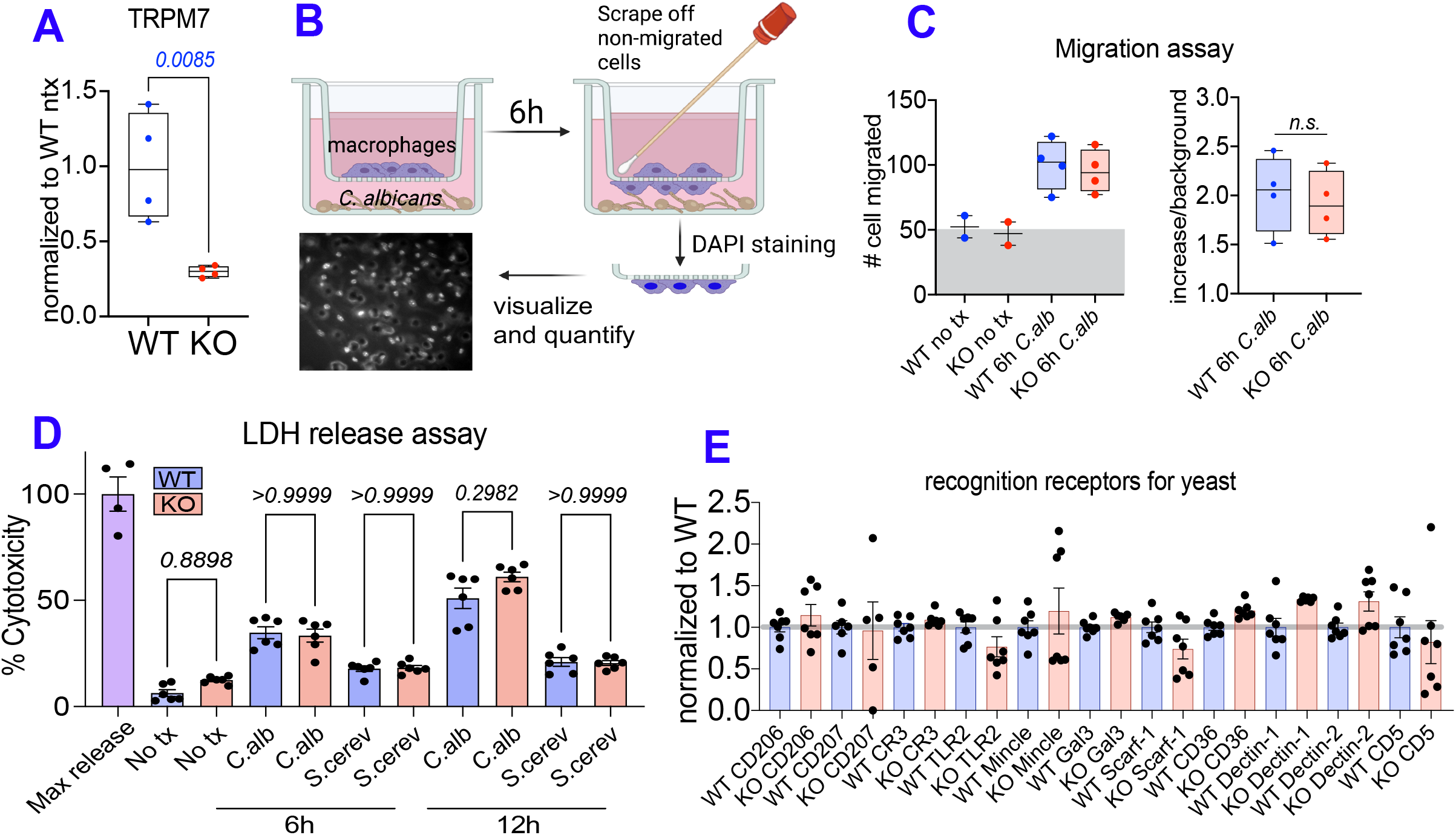
(A) TRPM7 gene expression analysis (qPCR) in primary macrophages, isolated from *TRPM7*^*fl/fl*^ (WT, blue dots) and *TRPM7*^*fl/fl*^ *LysM-Cre* (KO, red dots), normalized to WT macrophage expression. Statistics: two-tailed *t* test (Mann-Whitney). (B) Graphical outline of a transwell migration assay performed with primary macrophages. (C) Quantification of transwell migration assay. *Left panel*: the number of migrated WT (blue) and KO (red) macrophages at baseline (no tx, media only control) and 6h post-incubation with *C. albicans*. Each dot represents an average of 10 different ROIs. *Right panel*: data expressed as increased over baseline migration for WT (blue) and KO (red) macrophages incubated with *C. albicans* for 6h. Statistics: two-tailed *t* test. (D) Lactate dehydrogenase (LDH) release assay assessing macrophage viability during incubation with yeast species: *C. albicans* (*C. alb*) or *S. cerevisiae* (*S. cerev*) for 6h and 12h. WT macrophages (*n*=6, blue) and KO macrophages (*n*=6 red), positive control (*n*=4, purple). Statistics show ordinary one-way ANOVA with multiple comparisons. (E) Gene expression analysis (qPCR) of selected receptors at baseline in WT (blue) and TRPM7 KO (red) primary macrophages. Data are represented as fold change over WT (*n*=4-7, as indicated in the figure).

**Supplementary Figure 3.**
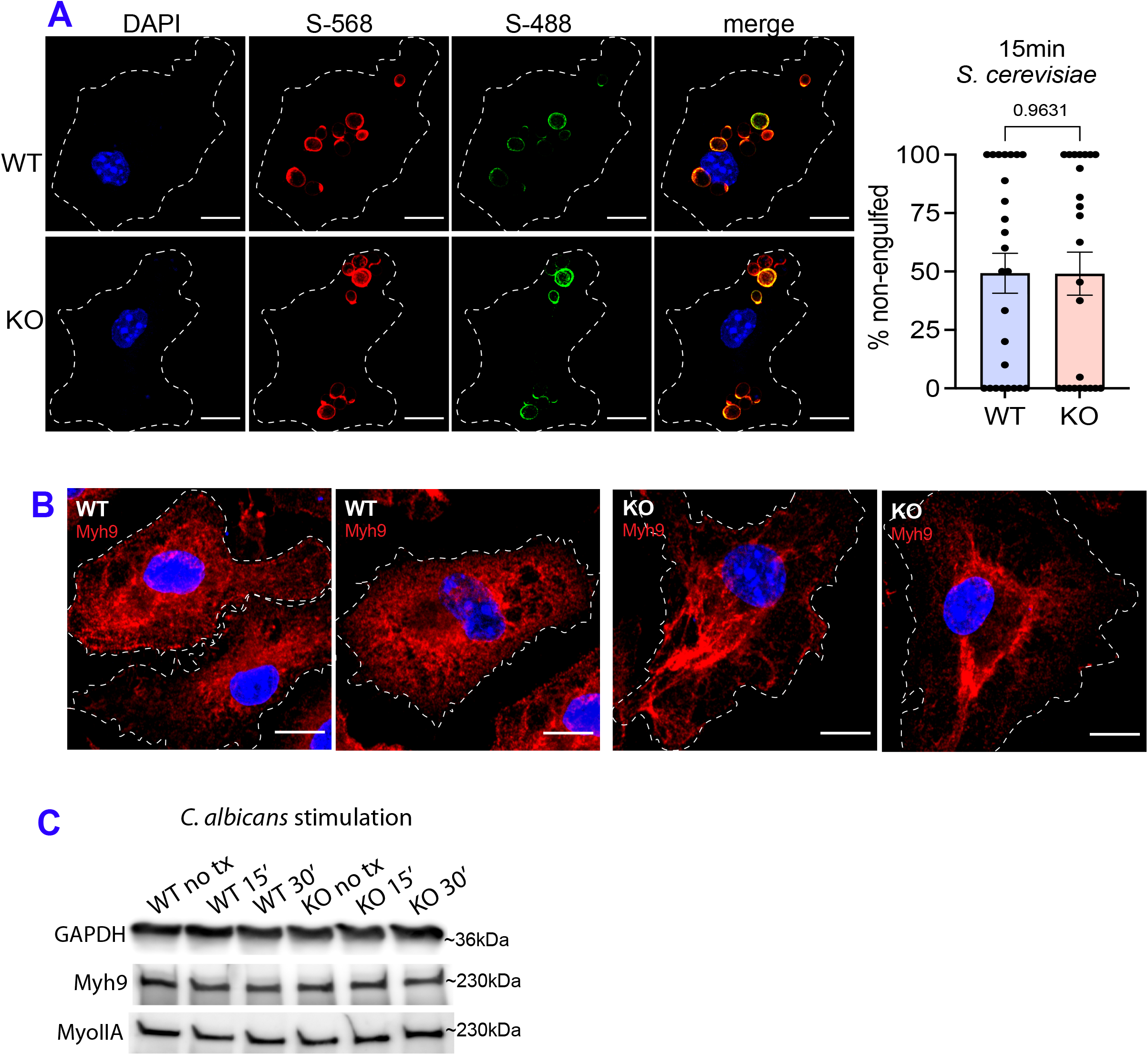
(A) Representative confocal images showing single channels and a merged image for WT (*upper panel)* and KO (*lower panel*) macrophages. *Right panel*: Quantification of the partially engulfed *S. cerevisiae* at 15 min for WT (blue) and KO (red) BMDMs. Error bars represent SEM. Statistics were performed with a two-tailed *t* test. Scale bar= 10 μm. (B) Representative maximal z-stack projection images of myosin IIA (Myh9, red) staining in WT and KO macrophages. DAPI stain in blue shows macrophage nuclei. Scale bar=10 μm. (C) Western blot of lysates from WT and KO macrophages stimulated with *C. albicans* for indicated times, as outlined in the figure. GAPDH was used as a loading control. The blot probed for Myosin IIA (Myh9) expression with two different antibodies.

**Supplementary Figure 4.**
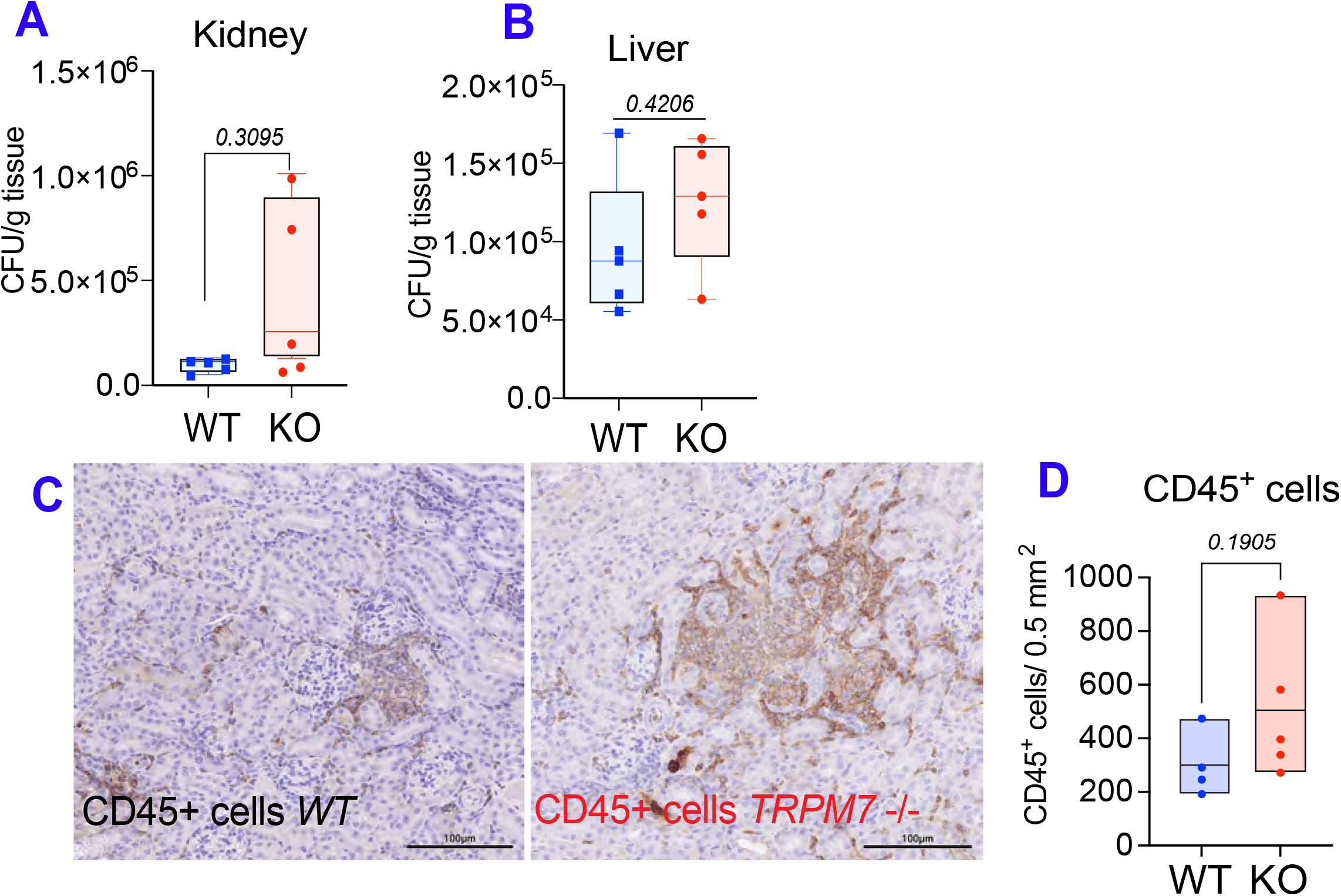
(A) Colony forming units (CFUs)/g tissue of *C. albicans* retrieved from kidneys at 48 h (*n*= 5 mice each, WT-blue, KO-red). Vehicle control mice had no CFUs. Statistics show two-tailed *t* test with SEM. (B) Colony forming units (CFUs)/ g tissue of *C. albicans* retrieved from livers at 48h (*n*= 5 mice each, WT-blue, KO-red). Vehicle control mice had no CFUs. Statistics show two-tailed *t* test with SEM. (C) Representative kidney sections stained with anti-CD45 antibody for WT (*left panel*) and KO (*right panel*). The scale bar is 100 μm. (D) Quantification of anti-CD45 stained kidney sections/0.5 mm^2^ area at 48 h post *C. albicans* injections in WT (blue, *n*=5) and KO (red, *n*= 5) mice. Data were plotted as floating bars and statistics were performed using two-tailed *t* test.

